# Structure of the mycobacterial ESX-5 Type VII Secretion System hexameric pore complex

**DOI:** 10.1101/2020.11.17.387225

**Authors:** Kathrine S. H. Beckham, Christina Ritter, Grzegorz Chojnowski, Edukondalu Mullapudi, Mandy Rettel, Mikhail M. Savitski, Simon A. Mortensen, Jan Kosinski, Matthias Wilmanns

## Abstract

To establish an infection, pathogenic mycobacteria use the Type VII secretion or ESX system to secrete virulence proteins across their cell envelope. The five ESX systems (ESX-1 to ESX-5) have evolved diverse functions in the cell, with the ESX-5 found almost exclusively in pathogens. Here we present a high-resolution cryo-electron microscopy structure of the hexameric ESX-5 Type VII secretion system. This 2.1 MDa membrane protein complex is built by a total of 30 subunits from six protomeric units, which are composed of the core components EccB_5_, EccC_5_, two copies of EccD_5_, and EccE_5_. The hexameric assembly of the overall ESX-5 complex is defined by specific inter-protomer interactions mediated by EccB_5_ and EccC_5_. The central transmembrane pore is formed by six pairs of EccC_5_ transmembrane helices that adopt a closed conformation in the absence of substrate in our structure. On the periplasmic face of the ESX-5 complex, we observe an extended arrangement of the six EccB_5_ subunits around a central cleft. Our structural findings provide molecular details of ESX-5 assembly and observations of the central secretion pore, which reveal insights into possible gating mechanisms used to regulate the transport of substrates.

## Main

Mycobacterial pathogens encode up to five Type VII secretion systems (ESX-1 to ESX-5), that are responsible for the secretion of a wide range of virulence proteins across the complex mycobacterial cell envelope^1^. The ESX-5 Type VII Secretion System (T7SS) is found almost exclusively in slow growing, pathogenic mycobacteria^2^ and has been shown to play a key role in nutrient uptake and immune modulation during an infection^3–5^. The importance of the ESX-5 and other ESX secretion systems to the virulence of mycobacterial pathogens makes these membrane spanning machineries an attractive target for the development of novel therapeutics^6^.

The first structure of a T7SS complex came from the *M. xenopi* ESX-5 system, which shares a high sequence similarity with the ESX-5 system from *M. tuberculosis*^7^. The low-resolution structure revealed a hexameric complex around a central pore with dimensions that suggested that it spans the inner mycobacterial membrane^7^. Recent high-resolution structures of the ESX-3 complex from *M. smegmatis* showed a dimeric assembly comprised of two EccB_3_:EccC_3_:EccD_3_:EccE_3_ protomer units with 1:1:2:1 stoichiometry^8,9^. This dimer was proposed to represent a building block of the hexameric holo-complex that would be required for secretion of substrates through the central pore. Docking of the dimeric ESX-3 structures into the ESX-5 low-resolution model^7^ did not provide, however, a satisfactory assembly of the pore due to the absence of EccC3 transmembrane helices and steric clashes in other parts of the model suggesting that significant structural reorganisation would be required^8,9^. Therefore, structural insights into the overall assembly of T7SS pore complex required for substrate translocation across the mycobacterial membrane have remained elusive, to date.

### Overall structural organisation of the hexameric ESX-5 pore complex

Here we used an integrative structural biology approach using single particle cryo-electron microscopy (cryo-EM) as primary source of structural information to elucidate the molecular basis of the T7SS hexameric pore architecture at high-resolution using the mycobacterial ESX-5 system as model. We expressed and purified the ESX-5 complex from *Mycobacterium xenopi* as previously described^7^ (Extended Data Figure 1).

An initial map without imposed symmetry at 3.9 Å resolution revealed substantial differences in the interpretability of different regions (Extended Data Figure 2). Consistent with the previous low-resolution model^7^, the membrane and cytoplasmic regions of the complex both display 6-fold symmetry, with the latter showing a larger extent of positional disorder. Based on this finding, we generated two maps for modelling these regions imposing C6 symmetry: a full map with a global resolution of 3.4 Å and a local refined map of the cytosolic part of a protomeric unit at 3.0 Å resolution (Figure 1a). In summary, the transmembrane domains and cytoplasmic domains close to the membrane of the ESX-5 complex could be built *de novo* to 88 % completeness with detectable sequence register (EccB_5_ 18-73; EccC_5_ 12-417; EccD_5_-1 23-502; EccD_5_-2 18-494; EccE_5_ 95-332) (Figure 1d, Extended Data Figure 4).

**Figure 1.**
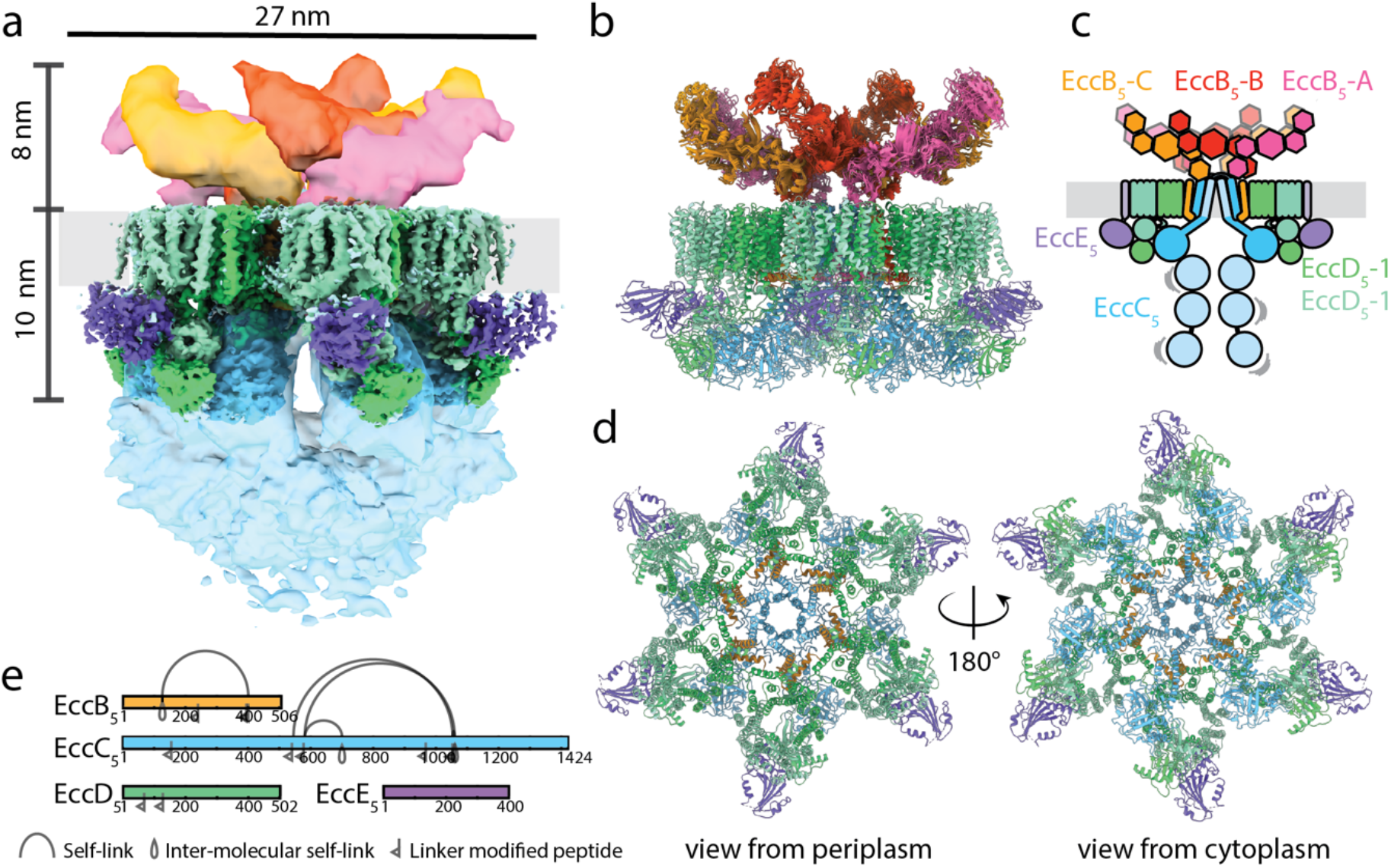
Overall architecture of the ESX-5 complex. **a**, Composite EM map of the ESX-5 complex. Periplasmic regions are coloured in yellow, orange, pink, transmembrane and membrane-proximal cytosolic region are coloured according to the scheme shown in **c** and the distal cytosolic segment tentatively corresponding to the ATPase domains is shown in blue. The membrane region is indicated in light grey. **b**, Side view of the complete atomic co-ordinate model of the membrane complex, including the periplasmic EccB_5_ region and the inner pore-forming EccC_5_ helical ring within the transmembrane segment, which have been modelled with ensemble approaches. **c**) Schematic of the ESX-5 complex showing the organisation of the components EccB_5_, EccC_5_, EccD_5_ and EccE_5_. The different EccB_5_ domains are coloured to show their location in the segmented EM map in **a**. Uninterpretable or low-resolution density has been shown as a lighter shade for EccC_5_ and EccE_5_. **d**, Top and bottom views of the rigid core of the ESX-5 membrane complex. **e**, Crosslinking pattern of the ESX-5 complex, crosslinks were observed in the EccB_5_ and EccC_5_ components.

In contrast to the 6-fold symmetry observed for the transmembrane and cytosolic segments of the ESX-5 secretion complex, the periplasmic region displays approximate C2 symmetry. Therefore, a third map without imposed symmetry constraints at 4.6 Å resolution was used for interpreting the periplasmic region of the complex. As the resolution of this map was not sufficient for *de novo* model building, we used a homology model of the periplasmic EccB_5_ domain^10^ and restraints for estimated residue pair distances from crosslinking analysis to build an integrative ensemble model comprising all six copies of EccB_5_ (Figure 1b,c Figure 2b-c) at an estimated precision of 7 Å (Methods, Extended Data Figure 5, Supplementary Table 2). This model provides insights into the arrangement of periplasmic EccB_5_ domains, which may act as a docking platform for additional components required to bridge the periplasm and reach the outer membrane. Taking the resulting models of the cytosolic, transmembrane and periplasmic segments together, we unravelled the architecture of the ESX-5 pore complex (Figure 1).

**Figure 2.**
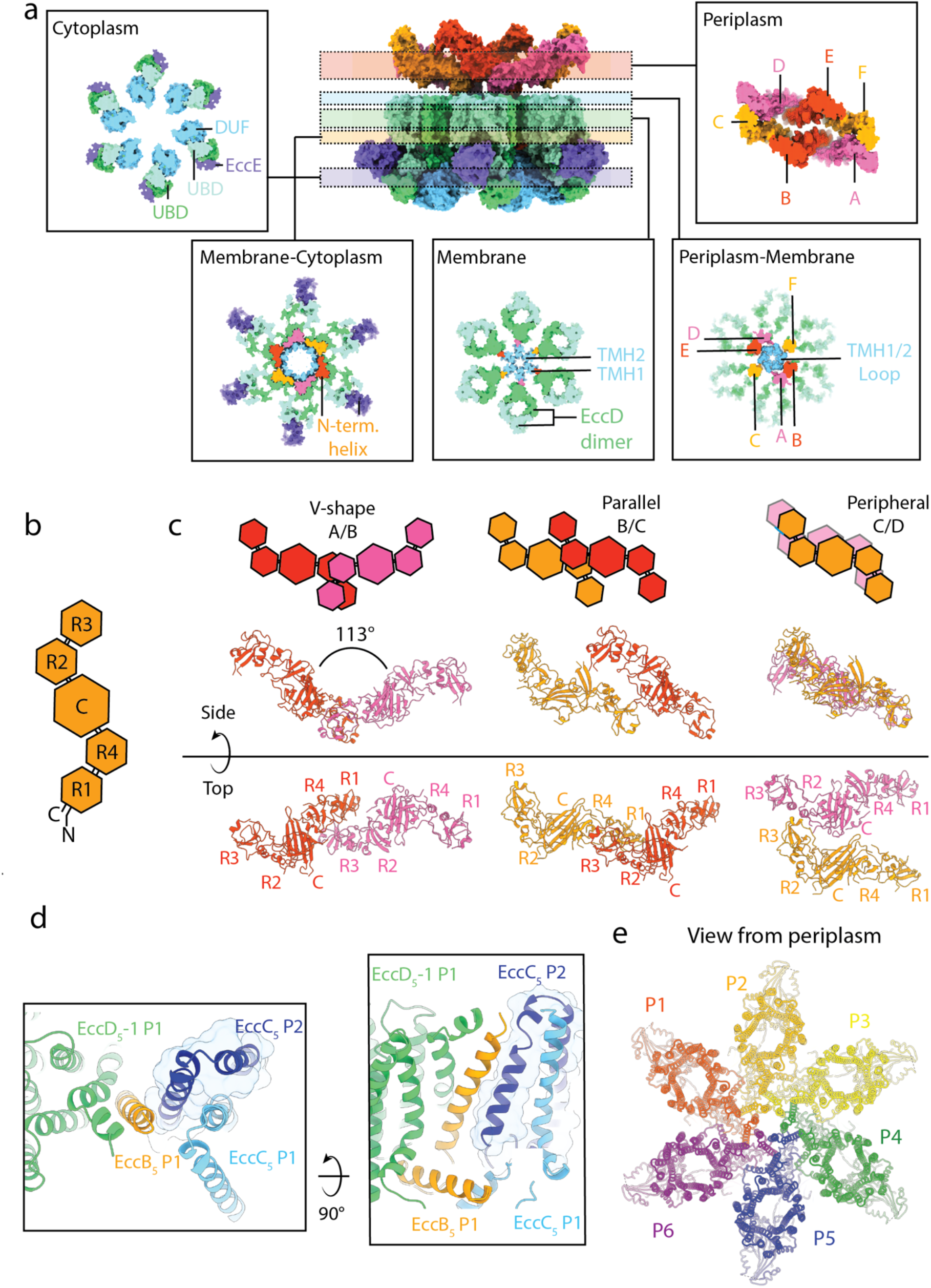
Subunit interactions and assemblies within the ESX-5 complex. **a**, Cross-sections of the ESX-5 complex, displayed as a surface representation of the atomic model shown as insets. Each inset highlights the subunit/subunit interactions occurring on the periplasmic, periplasmic-membrane, membrane, membrane-cytoplasmic and cytoplasmic level. Key features described in the text are highlighted **b**, Schematic representation of the periplasmic part of EccB_5_, indicating the repeat domains (R1-R4) and the central domain (C). **c**, Overview of three types of EccB_5_ dimers, shown in side and top view, annotated as ‘V-shape’, ‘parallel’ or ‘peripheral’. These three EccB_5_ dimers assemble into the characteristic dimer-of-trimer arrangement of the periplasmic face structure of the overall ESX-5 complex (see inlet of panel a), which does not follow the 6-fold symmetry of all other parts of the complex. In the top view, the subdomains in each EccB_5_ protomer are labelled, indicating specific EccB_5_ domain/domain involvement different for the three distinct EccB_5_ dimers observed. **d**, Inter-protomer interactions occurring between the EccC_5_ transmembrane helix 1 (TMD1) arising from protomer 1 (P1) and EccB TMD arising from protomer 2 (P2). The two panels represent the top and side view, respectively. **e**, Overview of domain swap interactions occurring between protomeric units when viewed from the periplasm, each protomer has a different colour.

### Architecture of the ESX-5 protomer

The ESX-5 complex is formed of six protomeric units comprised of EccB_5_, EccC_5_, two copies of EccD_5_ and EccE_5_. The core of the protomer is formed by an elliptical ring-shaped dimer of EccD_5_ comprised of 22 transmembrane helices (TMH). One of the EccD_5_ molecules is proximal and the second one distal to the central pore, annotated as EccD_5_-1 and EccD_5_-2, respectively. The interaction between EccD_5_-1 and EccD_5_-2 is mediated via hydrophobic interactions (Figure 2 Membrane inset, Extended Data Figure 3d) between TMH 9 and 10 from EccD_5_-1 and TMH 1 and 2 from EccD_5_-2 and *vice versa*.

The asymmetric ring-shape arrangement of EccD_5_-1 and EccD_5_-2 pairs enclose a cavity in the membrane that is partly filled with uninterpretable density, most likely attributable to additional lipids that were identified as part of our mass spectrometry analysis of the overall complex (Extended Data Figure 3d-e). The EccD_5_ ring-like arrangement is further supported by interactions at the cytoplasmic face formed by TMH6 and the elongated TMH7 that are connected via a long loop with unrelated conformations in each of the two EccD_5_ subunits. Due to the approximate two-fold symmetry of each EccD_5_ dimer, TMH11 of EccD_5_-2 is found most distal to the central pore whereas TMH11 of EccD_5_-1 is closest to the pore, both of them with a distinct diagonal orientation. TMH11 from EccD_5_-1 connects to the transmembrane helix of EccB_5_, thereby establishing a scaffold for the central pore. In a similar manner EccD_5_-2 TMH11 interacts with the transmembrane domain of EccE_5_, however this helix is not included in our high-resolution model due to weaker density, indicating flexibility at the periphery (Extended Data Figure 4). The central role of EccD_5_ as the scaffold for protomer organisation is further supported by studies where mutations in EccD led to a reduction in secretion of ESX and PE/PPE proteins^4,11^, likely due to disruption of key interactions stabilising the protomer.

EccD_5_ also acts as a key connector to the cytoplasmic regions of the ESX-5 protomer (Figure 2a Cytoplasm inset, Extended Data Figure 4a-d). The ubiquitin-like domains (UBDs) dimerise and interact with the cytoplasmic domain of EccE_5_ at the periphery of the complex. On the side proximal to the central pore, EccD_5_ UBDs interact with the EccC_5_ domain of unknown function (DUF), that follows the EccC_5_ stalk domain. The stalk domain is flanked by the long loop between TMH6 and TMH7 of EccD_5_-1 as well as the linker between EccD_5_-2 TMH1 and the EccD_5_-2 UBD. In addition to these interactions, at the interface between membrane and cytoplasm (Figure 2a membrane cytoplasm inset), we observe several specific electrostatic interactions of the stalk domain with the N-terminal EccB_5_ helix, situated parallel to the membrane. As for the membrane region, the EccD_5_ UBD dimer acts as a central scaffold for the other ESX-5 cytosolic subunits. The overall architecture of the ESX-5 protomer is similar to the ESX-3 protomer^8,9^, with key differences in the position of the EccC_5_ helices, described further below.

In our structure, the DUF domain represents the last rigid component of the ESX-5 complex. We attribute the less interpretable density for the remaining C-terminal part of EccC_5_ to the increased flexibility of ATPase domains 1 to 3, which has been previously observed^7–9^. However, at low threshold values the C1 map indicates the conformational space occupied by the ATPase domains (Extended Data Figure 6a). The flexibility of EccC appears to be a common feature to the ESX systems and was similarly observed in the ESX-3 dimer^8,9^, which may support the hypothesis that this flexibility is key to the function of the T7SS.

### EccB_5_ forms a diverse interaction network in the periplasm

The EccB_5_ model resulting from our integrative structural biology approach shows two distinct EccB_5_ trimers (EccB_5_-A-C and EccB_5_-D-E highlighted in Figure 2a Periplasm inset), forming an elongated keel-shaped assembly with overall dimensions of 20 nm in length, 10 nm in width and 8 nm in height divided by a central cleft. The dimer of trimers can be further subdivided into three distinct EccB_5_ dimers, denoted as ‘V-shape’, ‘Parallel’ and ‘Peripheral’ (Figure 2c). The angle between the two V-branches is established by interactions R1+R4 (EccB_5_-A) / R1+R4 (EccB_5_-B) and has a well-defined mean value of 113° (Extended Data Figure 7). Interestingly, the ‘V-shaped’ dimer conformation was also observed in the ESX-3 dimer^8,9^. EccB_5_-B is further involved in an interface with EccB_5_-C, where B and C are shifted with respect to each other by about 5 nm forming the ‘parallel dimer’, which is stabilised by interactions between the central (‘C’) domains: (EccB_5_-B) / R1+R4 (EccB_5_-C) and C (EccB_5_-C) / R2+R3 (EccB_5_-B). The ‘peripheral’ dimer forms the interface between the trimers along the long axis of the keel. These interactions are mediated by R2+R3 of EccB_5_-C from one trimer and EccB_5_-D (Figure 2c) from the other trimer. In the centre of the EccB_5_ assembly is a cleft region extending towards the central pore (Extended Data Figure 6b). The extensive interactions occurring within the periplasm between different protomers mediated by EccB_5_ suggest that the rearrangement of the EccB_5_ periplasmic domains following assembly of the six promoters is key to stabilising the hexameric holo-complex.

### Structural basis of hexameric ESX-5 assembly

The homology of EccC_5_ to FtsK AAA+ ATPases has led to suggestions the EccC_5_ may be the main driver of a hexameric assembly^12^. However, the stalk or DUF domains of EccC_5_ do not self-associate and owing to the flexibility of the EccC_5_ ATPase domains, it is unlikely that they are involved in permanent inter-EccC_5_ interactions. Together this suggests that other interactions are responsible to stabilising the T7SS in a hexameric assembly.

Besides the numerous inter-protomer interactions observed in the periplasm between EccB_5_, that were described above, a further key interaction for hexamerisation occurs between the transmembrane domains of EccB_5_ and EccC_5_. The EccC_5_ TMH1 of one protomer interacts with the EccB_5_ TMH of the neighbouring protomer via hydrophobic interactions and is further stabilised by the EccC_5_ TMH2 (Figure 2d). This interaction, which can mechanistically be described as domain swapping, is repeated in an anti-clockwise fashion when viewed from the periplasm (Figure 2e). This interaction acts to secure one protomer to the next by an interlocking mechanism in the membrane. Due to the low-resolution in this area for the ESX-3 dimer structure, this inter-protomer interaction was not observed^8,9^. Based on our model we envisage that within the membrane initial contact between protomers triggers the swap of the EccC_5_ helix to the next protomer. In turn, this may lead to changes in the orientation of the EccB_5_ periplasmic domains causing them to interlock, forming a stable unit in the periplasm. However, to confirm this model a structure of a monomeric protomer would be required to observe the pre-domain swap conformation.

### Insights into the central ESX-5 pore

The TM helices of EccC_5_ from each of the six protomers contribute to the formation of a central pore, a striking feature of the hexameric complex (Figure 3). The centre of the pore contains the TMH2 of EccC_5_, that interacts through helical bundle interactions with TMH1 of the neighbouring EccC_5_ subunit. In contrast to most other TM helices of the complex, the density of the TMH2 is less well defined (Figure 3c-d) which implies that this helix may be flexible and adopt a range of orientations, as indicated by our calculated ensemble model (Figure 3b). Central in these models is a highly conserved proline P73 (Supplementary Information 1, Figure 3a-c) triggering a significant kink in the middle of TMH2 and increased flexibility of its C-terminal part. As observed in other transport proteins, proline residues positioned in the middle of transmembrane helices may be key for their regulation and function^13^. For example, recent work describing a range of conformational states of an ABC exporter during its transport cycle, showed the role of a flexible helix with a central proline residue that acts as “gatekeeper”, regulating substrate access to the binding cavity^14^. We therefore suggest that P73 acts as a hinge point facilitating conformational flexibility of the pore, required for substrate transport.

**Figure 3.**
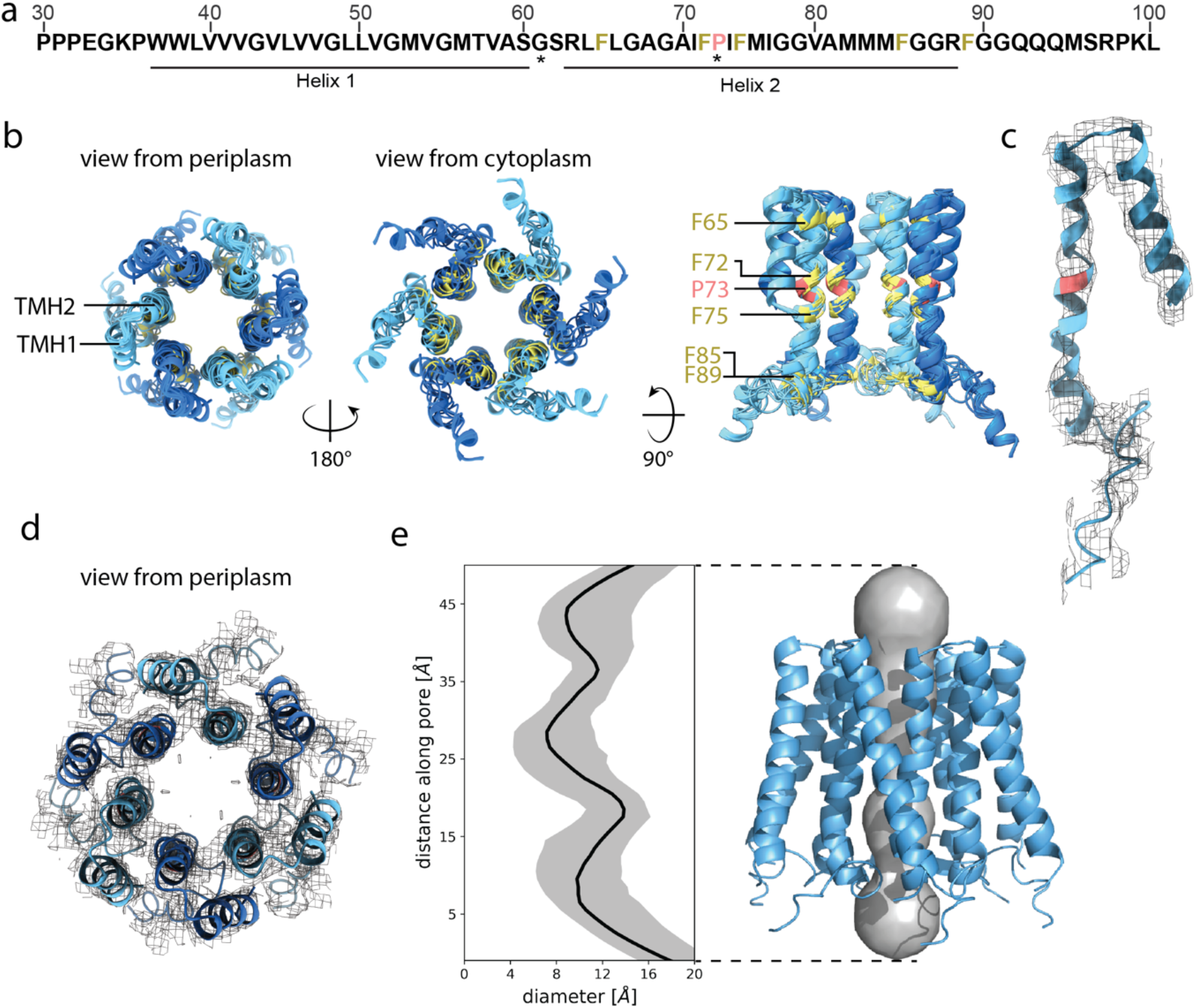
Central pore of the ESX-5 secretion system. **a**, *M. xenopi* EccC_5_ sequence near the N-terminus (residues 30-100) highlighting the two transmembrane helices observed in the structure of the overall ESX-5 complex (for further details see Supplementary Information 1). Invariant residues are indicated by asterisk. Colours correspond to the model shown in **b**. **b**, Overlay of the top ten highest scoring Rosetta^14^ models viewed from the periplasm, the cytoplasm and along the membrane. Conserved residues have been coloured (proline, red; phenylalanine, yellow). **c**, EM density of the full C6 map around TMH2 of the top scoring Rosetta model. **d**, View from the periplasm of the EM density corresponding to the central pore. The top scoring Rosetta model is shown **e**, Analysis of the pore diameter from 100 Rosetta models with HOLE^39^, median pore diameter (black) and 90% confidence interval (grey). Side view of EccC_5_ pore helices of a top scoring model showing the pore diameter analysis.

TMH2 also comprises a number of phenylalanine and methionine residues that are common to this helix across different ESX systems (Figure 3a, Supplementary Table 1). In the ESX-5 structure P73 is flanked by F66, F72 and F75, inevitably orienting some of these aromatic side chains towards the inner surface of the pore and thus reducing the pore diameter. To quantify this, we analysed the diameter of the 100 highest-scoring ensemble models to sample the range of potential conformations of TMH2 (Figure 3e). These results suggest that the central pore, in this structure, may have up to three narrow constrictions reducing the pore diameter to less than 1.0 nm. As this diameter is too small to secrete folded heterodimers with a size of around 2.2 nm^15^ through the pore, we believe that our model in the absence of bound substrate represents a closed state of the secretion system.

Analysis of the central pore of the ESX-5 system has shown intriguing similarities to other transport systems. The lining of secretion pores with bulky hydrophobic amino acids has been reported for other secretion machineries, which act to gate the pore, regulating secretion. For example, in A/B type toxins, such as the anthrax toxin from *Bacillus anthracis*, the secretion of the lethal factor through the protective antigen membrane spanning complex is modulated by phenylalanine rings or ‘⌽ clamp’ at the top of channel which constrict the channel width to approximately 6 Å, thereby restricting the passage of a folded lethal factor^16,17^. Another example comes from the export apparatus complex of T3SS where a highly conserved Met-Met-Met loop forms a molecular gasket constructing the channel to less than 10 Å and thereby prohibiting secretion^18^. These systems do not share any sequence similarity with the ESX system, suggesting that the “default” closed state of the pore is a fundamental principle of bacterial secretion machineries, evolving through convergent mechanisms on different transmembrane protein scaffolds.

### Discussion and Outlook

Our cryo-EM structure of the hexameric *M. xenopi* ESX-5 pore complex provides insights into the overall architecture of the T7SS secretion system. The high-resolution structure of the membrane regions reveals key interactions within and between protomers explaining how the hexamer is formed. The structural analysis of the pore of the ESX complex, which in our structure is in a closed state, hints to how this pore is gated. Future work to delineate the steps leading to pore opening upon the binding of secreted substrates will provide key mechanistic insights into how transport is regulated. On the protomer level, the structural conservation between the ESX systems is high, however the propensity of the different systems to form a hexamer may differ, which in turn could further implicate the role of external factors to trigger hexamerisation in the different systems. For future drug discovery endeavours, targeting different assembly states of the T7SS may prove an efficient strategy for inhibiting secretion of key virulence proteins.

## Methods

### Molecular Biology

Polymerase chain reaction (PCR) was performed using Q5 DNA polymerase (New England Biolabs). For cloning, *E. coli* DH5α was used. The *eccD_5_* gene was amplified by PCR to include the N-terminal ubiquitin like domain (residues 1–129) and inserted into the pMyNT vector using SliCE methods^19,20^, generating pMyNT-EccD_5_^129^ The *M. xenopi* ESX-5 complex was expressed from pMV-ESX-5 vector in *M. smegmatis* as previously described^7^.

### Protein expression purification

Expression vectors were transformed into *M. smegmatis* mc^2^155 *groEL1ΔC*^21^ and grown in Middlebrook 7H9 medium (BD Biosciences) supplemented with 0.2 % (w/v) glucose (Carl Roth), 0.05% (v/v) Tween-80 (Carl Roth) and 0.2% (v/v) glycerol (Carl Roth) with appropriate antibiotics. For expression of the ESX-5 complex cells were cultured to an optical density (OD) at 600 nm of 1.5 and pelleted by centrifugation. For the production of EccD_5_^129^ from an acetamidase inducible promoter cells were grown to and OD 600 nm of 1.0 and induced with 1 % acetamide and cultured for a further 24 h at 37 °C and pelleted by centrifugation. The ESX-5 complex was purified as previously described in Beckham et al, 2017^7^.

For the purification of EccD_5_^129^ cells were resuspended in buffer A (20 mM Tris pH 8.0, 300 mM NaCl, 20 mM imidazole) with EDTA-free protease inhibitors (Roche) and DNase (Sigma). Cells were lysed by high-pressure emulsification, and unbroken cells were removed by centrifugation at 4 °C for 20 min (19,000g).

### Crosslinking mass spectrometry analysis

Purified complex at concentration of 50 μg (1 mg/ml) was crosslinked by addition of an iso-stoichiometric mixture of H12/D12 isotope-coded, di-succinimidyl-suberate (DSS, Creative Molecules). The crosslinking reaction (final concentration 1mM) was incubated for 30 min at 37°C and quenched by addition of ammonium bicarbonate to a final concentration of 50 mM for 10 min at 37°C. Crosslinked proteins were denatured using urea and RapiGest (Waters) at a final concentration of 4 M and 0.05% (w/v), respectively. Samples were reduced using 10 mM dithiothreitol (30 min at 37°C), and cysteines were carbamidomethylated with 15 mM iodoacetamide (30 min in the dark). Protein digestion was performed using 1:100 (w/w) LysC (Wako Chemicals) for 4 h at 37°C and then finalised with 1:50 (w/w) trypsin (Promega) overnight at 37°C, after the urea concentration was diluted to 1.5 M. Samples were then acidified with 10% (v/v) TFA and desalted using OASIS^®^ HLB μElution Plate (Waters). Crosslinked peptides were enriched using size exclusion chromatography^22^.

Collected SEC fractions were analysed by liquid chromatography-coupled tandem mass spectrometry (MS/MS) using a nanoAcquity Ultra Performance Liquid Chromatography (UPLC) system (Waters) connected online to Linear Ion Trap Quadrupole (LTQ)-Orbitrap Velos Pro instrument (Thermo). Peptides were separated on a BEH300 C18 (75 × 250 mm, 1.7 mm) nanoAcquity UPLC column (Waters) using a stepwise 60-min gradient between 3% and 85% (v/v) acetonitrile in 0.1% (v/v) fusaric acid. Data acquisition was performed using a top-20 strategy where survey MS scans (m/z range 375–1,600) were acquired in the Orbitrap (R = 30,000) and up to 20 of the most abundant ions per full scan were fragmented by collision-induced dissociation (normalized collision energy = 40, activation Q = 0.250) and analysed in the LTQ. In order to focus the acquisition on larger crosslinked peptides, charge states 1, 2 and unknown were rejected. Dynamic exclusion was enabled with repeat count = 1, exclusion duration = 60 s, list size = 500, and mass window ± 15 ppm. Ion target values were 1,000,000 (or 500 ms maximum fill time) for full scans and 10,000 (or 50 ms maximum fill time) for MS/MS scans. The sample was analysed in technical duplicates.

To assign the fragment ion spectra, raw files were converted to centroid mzXML format using a raw converter and then searched using xQuest^23^ against a FASTA database containing the sequences of the crosslinked proteins. Posterior probabilities were calculated using xProphet^23^, and results were filtered using the following parameters: false discovery rate = 0.05, min Δscore = 0.95, MS1 tolerance window of −4 to + 7 ppm, identity (Id) score > ^36^.

### X-ray crystallography and data processing

EccD_5_^129^ crystallised in initial conditions from the Morpheus screen (Molecular Dimensions) containing 0.06 M MgCl_2_, CaCl_2_, 0.1 M Tris:bicine pH 8.5, 10 % OEG 20k, 20 % PEG MME 550. Diffraction data were collected at EMBL beamline P13 at the PETRA III storage ring (DESY, Hamburg, Germany). The data were processed with XDS^24^ and merged with AIMLESS^24,25^ and the relevant statistics are shown in Table S1. We used the EccD1129 model from *M. tuberculosis* (PDB: 4KV2) as a molecular replacement candidate (45% sequence identity to *M. xenopi* EccD_5_). After the successful placement of the model using Phaser^26^, manual building was performed in Coot^27^. The model was refined using REFMAC5^28^.

### Cryo EM Sample preparation and data acquisition

For cryo-EM, 3.6 μl of the ESX-5 void peak fraction was applied on freshly glow-discharged Quantifoil R2/1 Cu 200 mesh grids with 2 nm continuous carbon. The sample was blotted for 2 s and vitrified in a liquid propane/ethane mix using a Vitrobot Mark IV at 10 °C and 100% humidity. The grid was screened at the cryo-EM facility at CSSB (Hamburg, Germany) and high-resolution cryo-EM data were collected on a Titan Krios operated at 300 kV (Thermo Fisher Scientific FEI) equipped with a K3 direct detection camera (Gatan) and a BioQuantum K3 energy filter (Gatan) operated by SerialEM^29^ at the EMBL Cryo-Electron Microscopy Service Platform (Heidelberg, Germany). A total of 27.873 movies with 40 frames were recorded in counting mode, with a total dose of 49.34 e/Å^2^ and a pixel size of 0.645 Å. The underfocus range was set to 0.7 μm – 1.7 μm with a step size of 0.1 μm.

### Data processing

Data processing was performed in cryoSPARC^30^ and is visualised in Extended Data Figure 2. First, movie frames were aligned and local motion was corrected for using patch-motion correction. The contrast transfer function (CTF)-landscape of each micrograph was estimated using patch CTF-estimation. The exposures were curated based on local-motion distances and CTF-fit parameters. Particles were picked on the remaining 18,598 micrographs with a template-based particle picker. The templates were generated beforehand, based on a map obtained from an initial dataset. The picked particles were inspected and 635,219 were selected and subsequently extracted, using a box size of 58 nm. For 2D classification, the data was binned four times. Four rounds of 2D classification were performed. After each round, particles leading to intact classes were selected and included in the next round. A total of 284,402 particles passing these iterations were used to generate three *ab-initio* models. Particles corresponding to one class displaying a clear hexamer were further sorted with another round of 2D classification. The remaining selected 121,974 particles were reextracted and binned two times. An *ab-initio* model was generated and the data was refined using the non-uniform refinement algorithm in the absence of any imposed symmetry (C1). The map and initial model building attempts revealed six-fold symmetry for the transmembrane and membrane proximal-cytoplasmic regions. Based on this observation we generated another map, by imposing C6 symmetry. The resulting map at a global resolution of 3.4 Å revealed highest resolution in the transmembrane regions (approximately 2.8 Å – 3.5 Å) and lower resolution in the cytosolic part (approximately 4.0 Å - 5.5 Å).

To improve the resolution of the cytosolic part of each protomer, symmetry expansion was carried out followed by local refinement. The applied masks were created in UCSF Chimera^31^ and processed in cryoSPARC^30^. The generated map showed almost uniform distributed resolution for the cytoplasmic regions of about 3.0 Å. As no particle subtraction was performed beforehand, transmembrane helices of the protomer are still visible in the map.

Heterogenous refinement with three classes was further performed on the 121,974 particles to investigate for different conformations of the periplasmic region. One class was chosen to assess the 3D variability^30^ of cytoplasmic and periplasmic domains to further reduce heterogeneity. Based on the results three clusters could be identified. One of them showed a keel conformation in the periplasmic region and was further refined without and with C2 symmetry. The two other clusters remained heterogenous likely due to the misalignment of different orientations. To improve the alignment, the low pass filtered keel shape map (from cluster 1) was further refined and used as initial model for the refinement of the 121,974 particles. To achieve an improved separation of the keel shape another round of heterogenous refinement was carried out with subsequent refinement of one class without imposed symmetry. The resulting map had a global resolution of 4.6 Å, however the resolution of the periplasmic region is less then 10 Å.

### Atomic model building and refinement

As there are no reliable, high-resolution structures of any of the ESX-5 components or their homologues available in PDB^32^ we built *de novo* a model of the transmembrane and nearby cytoplasmic regions of the complex (EccB_5_ 18-73; EccC_5_ 12-417; EccD_5_-1 23-502; EccD_5_-2 18-494; EccE_5_ 95-332). An initial model was traced into a masked, focused-refinement map using ARP/wARP cryo-EM module with default parameters^33^. Next, domains for which we solved the high-resolution crystal structure (EccD_5_, residues 17:107, Extended Data Figure 3a-c) were fitted into the focused-refinement map as rigid-bodies using a Jiggle Fit tool from COOT27 (Extended Data Figure 6). The resulting model was completed manually using COOT in regions with local resolution allowing for unambiguous *de novo* model tracing. The interpretation of poorly resolved map regions was aided by alternative blurring and sharpening of the map in COOT. We used an iterative approach where each manual model building step was followed with sequence assignment using findMySequence program (Chojnowski, Simpkin, Keegan, and Rigden, unpublished), which allowed for an identification and correction of tracing errors (insertions, deletions). Loops that were resolved in the density, but difficult to trace manually, were built using the RosettaES density-guided enumerative-sampling algorithm from the Rosetta suite^34^. The complete protomer model built into a focused-refinement map was expanded to a complex using symmetry operations derived directly from the C6 symmetrised map using phenix.find_ncs_from_density^35^ and completed manually in COOT. Apart from solving minor symmetry conflicts we traced the model fragments that were resolved in the symmetrised map only. These included two transmembrane helices of EccC_5_ (TMH1 and TMH2; residues 37 to 94). Firstly, a better resolved in the density TMH1 was build *de novo* in COOT and assigned to the sequence using findMySequence program, which allowed for an unambiguous determination of the helix direction and sequence register. Subsequently, the second TM helix (TMH2) was built using the RosettaES density-guided enumerative-sampling algorithm followed with refinement with C6 symmetry. We also added to the model the most distant to the central pore EccD_5_-2 helix (TMH11) based on a model of the corresponding helix in EccD_5_-1. Finally, models of the protomer and full complex were refined against corresponding maps using phenix.real_space_refine^36^ with default parameters.

### Pore dimension analysis

The local-resolution of the central pore map region didn’t allow the determination side-chain conformations in the second EccC_5_ TM helix of (TMH2) and reliable measurement the pore dimensions. Therefore, we analysed a whole ensemble of tentative models resulting from a density-guided enumerative sampling refinement with C6 symmetry restraints implemented in Rosetta. The pore profiles of 100 lowest-energy models were calculated using HOLE program^37^.

### Integrative modelling

The model of the hexameric assembly of the periplasmic domains of EccB_5_ (aa. 74-490) was built using an integrative modelling protocol similar to the protocol used by us^38–40^. The modelling procedure described in more detail below is implemented as a custom software based on Integrative Modeling Platform (IMP)^41^ version 2.13 and Python Modeling Interface (PMI)^42^. All additional code and input files necessary to reproduce the steps will be released on Zenodo repository upon publication.

The transmembrane region of the ESX-5 structure built *de novo* as above and a homology model of the monomeric EccB_5_ periplasmic domain were used as input for modelling. The homology model of EccB_5_ was built using Modeller^42^ based on the crystal structure of EccB1 of *M. tuberculosis* (PDB ID: 3X3M10) and using the sequence alignment obtained from the HHpred server^43^. The non-symmetrised (C1) EM map and available EccB_5_ crosslinks were used as modelling restraints. Owing to the low-resolution (< 10 Å) of the periplasmic region, the high-frequency noise in the EM map was removed using a Gaussian filter with a standard deviation of 3 Å. Additionally, to limit the conformational space the fitting was performed using only a segment of the EM map not yet occupied by the transmembrane region of the ESX-5 structure. The models were additionally restrained using high-confidence crosslinks above ld-score 36 were. At this threshold, two crosslinks could be mapped to the EccB_5_ sequence and used for modelling: Lys125-Lys125, representing a self-assembly link from two different EccB_5_ copies, and Lys125-Lys400.

As the first step of the modelling, a large library of alternative fits to the EM map of the monomeric EccB_5_ structure was generated using the FitMap tool of the UCSF Chimera^34^. The fitting was performed using 100,000 random initial placements, cross-correlation about the mean as the fitting score (Chimera’s ‘cam’ score^31^, equivalent to Pearson correlation coefficient), and the requirement of at least 80% of the input structure being covered by the EM map envelope defined at a permissive density threshold. This resulted in unique alternative 9268 fits, after clustering.

Second, the resulting alternative fits of the monomeric EccB_5_ and the transmembrane region of the ESX-5 structure built *de novo* as above were used as input for the simultaneous fitting of six copies of EccB_5_ using the EM map and crosslink restraints. The fitting was performed through simulated annealing Monte Carlo optimisation that generates alternative configurations of the fits pre-calculated as above. The optimisation was performed independently 4,000 times with 12,000,000 Monte Carlo steps for each run. The sampling exhaustiveness was assessed by ensuring that i) score converges in individual runs, ii) no new better scoring models appear with extra runs, iii) score distributions in two random samples of the models are statistically similar (Extended Data Figure 6a-c). The scoring function for the optimisation was a sum of the EM fit restraint represented as the p-values of the precalculated domain fits (calculated as described in^38–40^), crosslinking restraints, clash score, connectivity distance between each EccB_5_ transmembrane helical segment and periplasmic domain next in sequence, a term preventing overlap of the protein mass with the transmembrane region, and a 2-fold symmetry restraint. During the optimisation, the structures were simultaneously represented at two resolutions: in Cα-atom representation and a coarse-grained representation, in which each 10-residue stretch was converted into one bead. The 10-residue bead representation was used for all restraints to increase computational efficiency except for the domain connectivity and crosslink restraints, for which the Cα-only representation was used for reasons of accuracy.

Finally, top scoring models from the previous step were subjected to a refinement coupled to an analysis of exhaustiveness of conformational sampling and estimation of model precision using a procedure proposed by Viswanath *et al*.^44^. To this end, the models from the first modelling stage (simultaneous fitting based on the alternative fits) were split into two random subsets. Top 30 models from each subset were refined using a Monte Carlo simulated annealing optimisation in which the structures were moved in the EM map with small rotational and translational increments. The scoring function consisted of cross-correlation to the EM map, domain connectivity restraint, clash score, a term preventing overlap of the protein mass with the transmembrane region, and a 2-fold symmetry restraint. Each of the 30 models was refined with 200 independent runs with 260,000 steps. Top scoring models from each of the two runs were selected leading to two independent samples of refined models (about 1000 models in each sample). The scores of the two samples were compared to each other to ensure convergence (Extended Data Figure 5d,e). The highest sampling precision at which sampling was exhaustive was determined based on the RMSD comparisons between all models and clustering at incremental RMSD thresholds using the statistical tests provided by Viswanath et al. (Extended Data Figure 5f). The two samples were then clustered at the resulting precision level (Extended Figure 6g) and for each cluster the model precision, defined as the average RMSD distance to cluster centroid, was calculated. The top ten scoring models from all refined models were taken as the final ensemble model of the ESX-5 with the EccB (Extended Data Figure 5h). All the top ten models satisfied both EccB crosslink restraints (with distance threshold of 30 Å, Extended Data Figure 5i).

The models will be deposited in the PDB-dev database upon publication.

## Acknowledgements

We thank Felix Weis and Wim Hagen for maintaining the EMBL cryo-EM facility in Heidelberg and Felix Weis for help with the data collection. We acknowledge the EM facility at the Centre for Structure Systems Biology, Hamburg. We would like to thank Annabel Parret and Luciano Ciccarelli for their previous work on the project. The synchrotron MX data was collected at beamline P13 operated by EMBL Hamburg at the PETRA III storage ring (DESY, Hamburg, Germany). We would like to thank Guillaume Pompidor for the assistance in using the beamline and Vasileios Rantos for help with scripts for exhaustiveness analysis of integrative modelling. This work was in part supported by a Joachim Herz Stiftung “Add-On Fellowship for Interdisciplinary Science” (awarded to K. S. H. B.) and by the Joachim-Herz-Stiftung Hamburg via the project Infectophysics.

## Author Contributions

M.W. supervised and supported the project. K.S.H.B produced samples of the *M. xenopi* ESX-5 complex suitable for crosslinking mass spectrometry and structural analysis, including biophysical characterisation and X-ray crystallography. S.A.M. performed high-resolution cryo-electron microscopy experiments; C.R. processed and interpreted the data, with support by E.M. K.S.H.B and M.R. conducted crosslinking mass spectrometry experiments M.R., supervised by M.M.S. G.C. conducted and developed approaches for rapid structural model building and interpretation. JK performed integrative modelling. K.S.H.B., C.R., G.C., J.K, and M.W. wrote the paper.

## Figures and Figure Legends

**Extended Data Figure 1.**
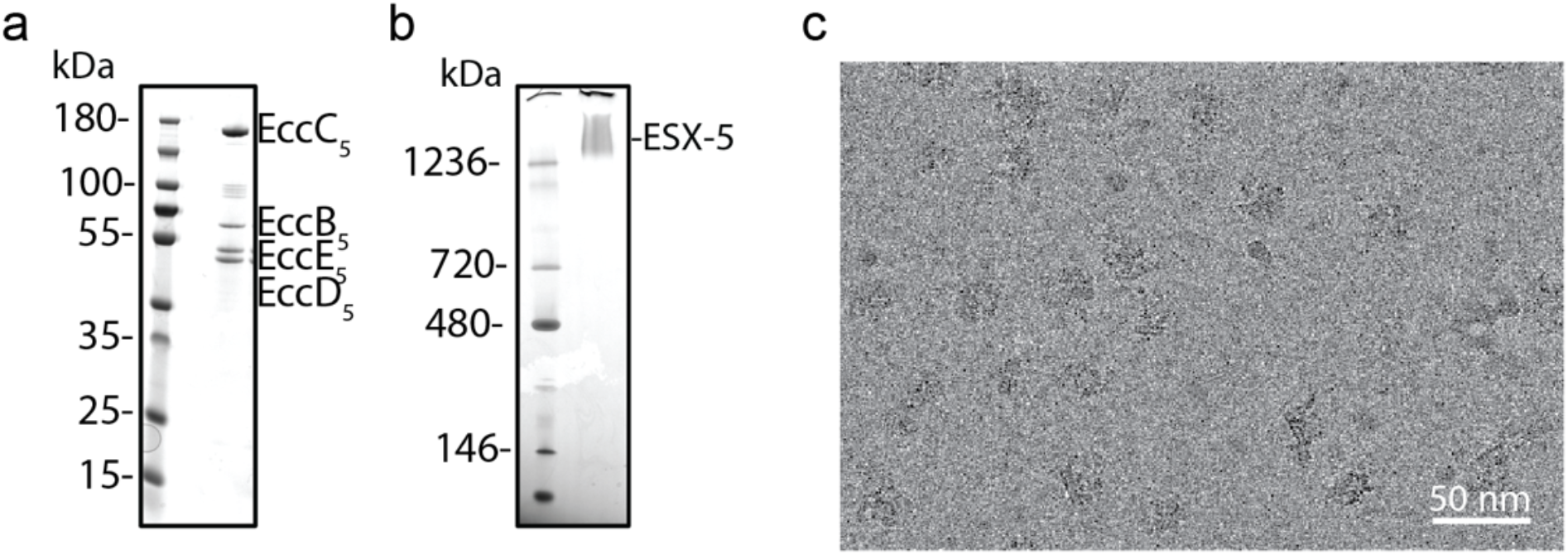
Purification of the ESX-5 complex. **a**, Coomassie stained SDS-PAGE and blue native PAGE gel (**b**) of purified ESX-5 complex used for cryo EM. **c**, Representative cryo electron micrograph showing ESX-5 particles. Scale bar, 50 nm.

**Extended Data Figure 2.**
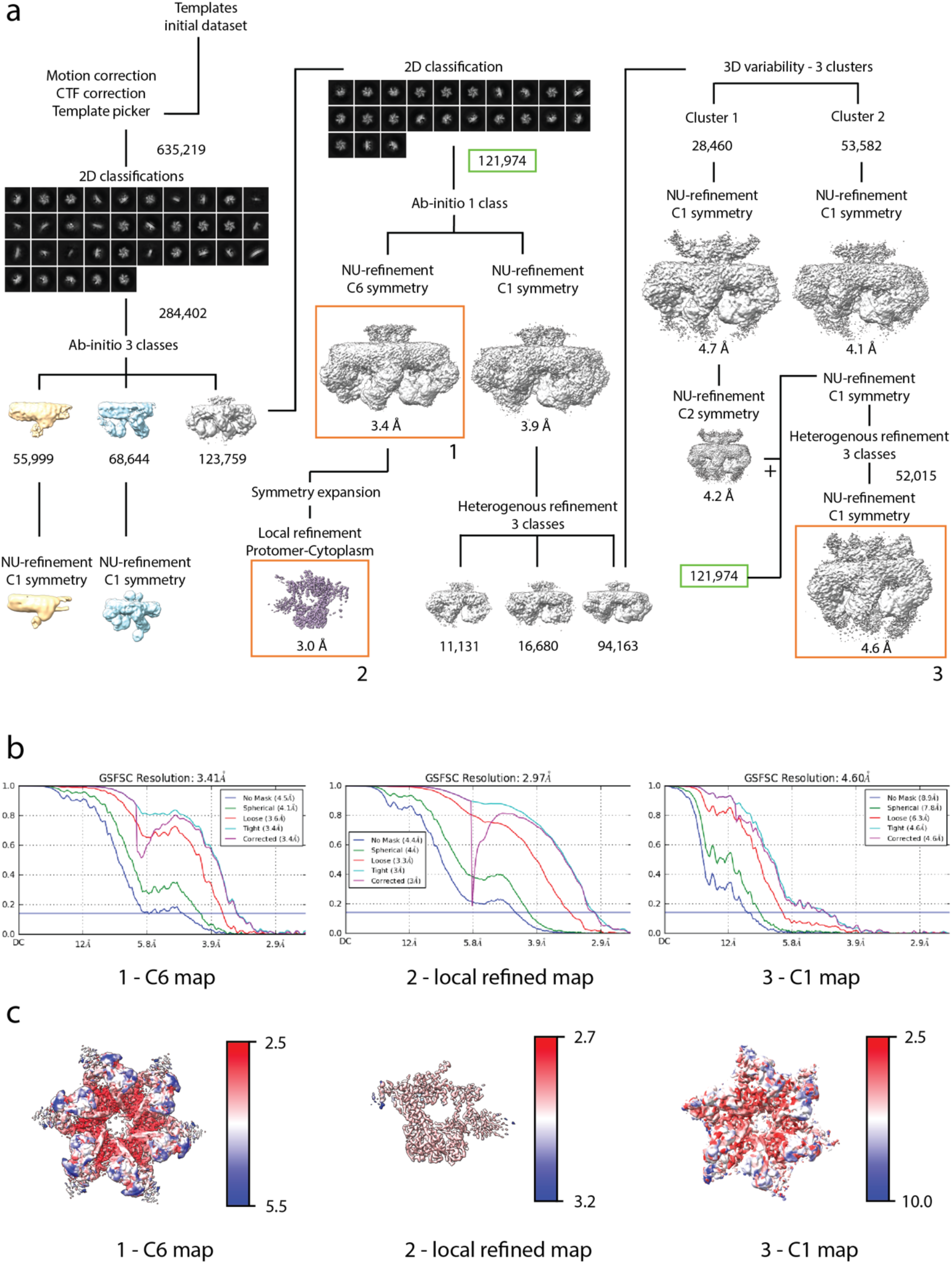
Cryo EM data processing. **a**, Cryo-EM image processing strategy applied to the 27.873 movies in cryoSPARC to generate a C6 symmetry map of the ESX-5 core at 3.4 A, a local refined map of a protomer at 3.0 A and a C1 map with low resolution features in the periplasm and cytoplasm (highlighted in orange and numbered 1 - 3). For each of the three maps **b**, Gold-standard Fourier shell correlation (GSFSC) curves with different masks calculated by cryoSPARC with resolution cutoff = 0.143 and **c**) local resolution estimations are shown.

**Extended Data Figure 3.**
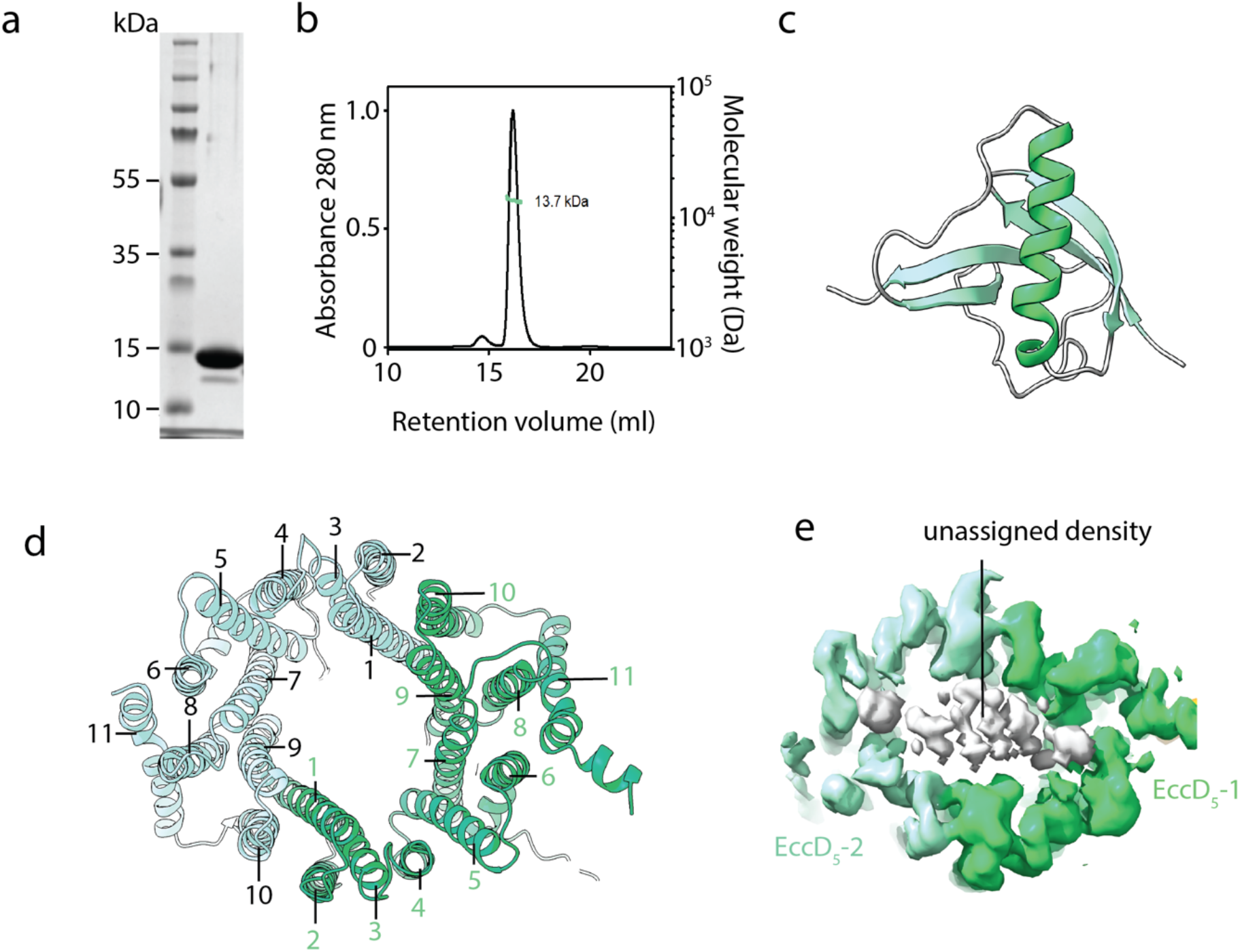
Structural characterisation of EccD_5_. **a**, SDS-PAGE showing the purified EccD_5_ ubiquitin like domain (EccD_5_^129^). **b**, SEC-MALS analysis of EccD_5_^129^, indicating a size of 13.7 kDa. **c**, Crystal structure of EccD_5_^129^ with secondary structure elements highlighted (helix, green; B-sheet, light green). **d**, View from the periplasm on the cavity formed by the transmembrane domains of the two EccD_5_ molecules. Helix numbers are indicated for EccD_5_-1 in green and for EccD_5_-2 in black **e**, View from the periplasm of the EM density of the EccD_5_ (green) transmembrane domains showing unassigned density (white).

**Extended Data Figure 4.**
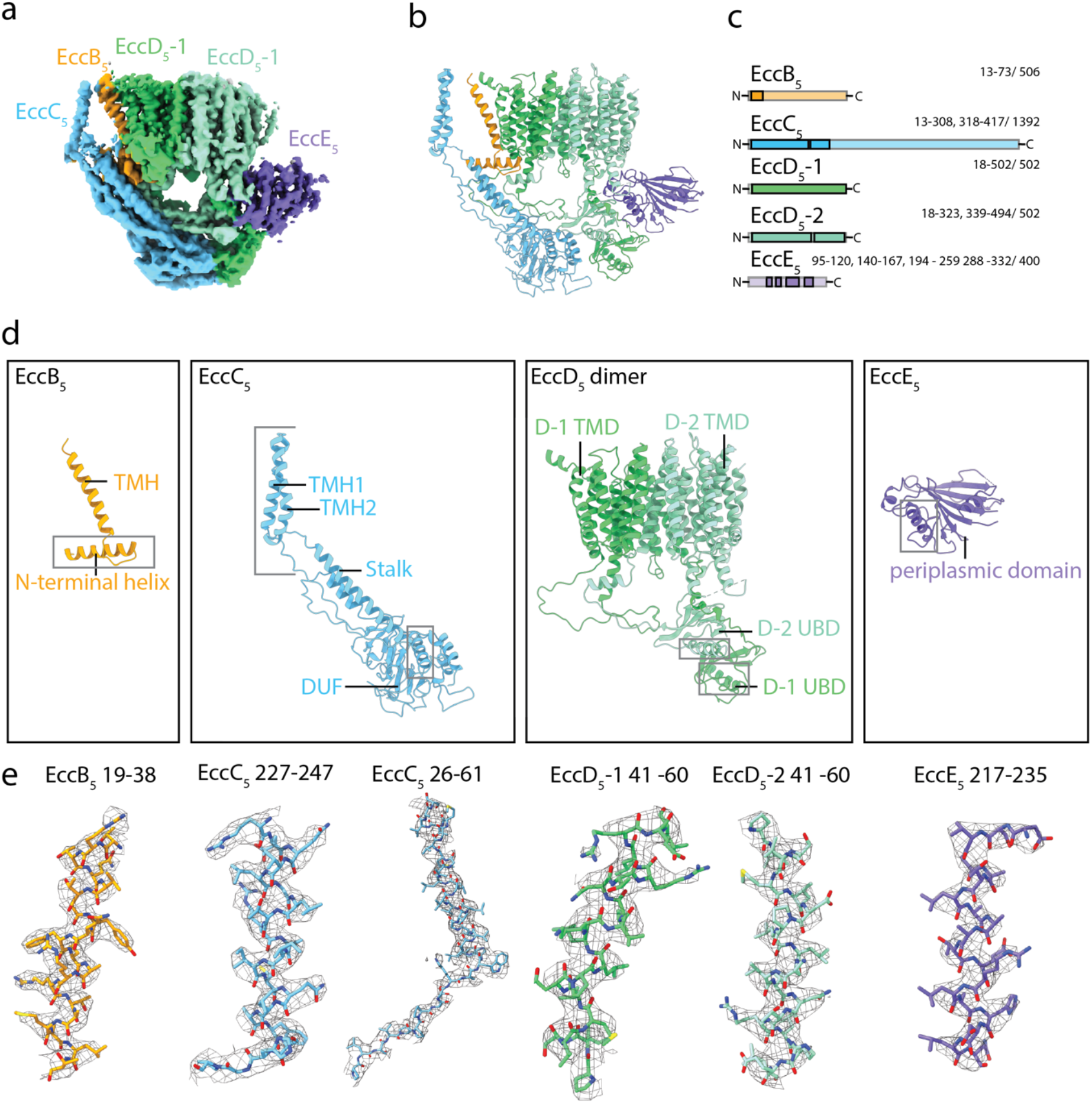
Overview of the ESX-5 protomer. **a**, EM map segment, showing a side view of the protomer. **b**, High-resolution model of a protomer shown from the side. **c**, Coverage of the high-resolution model with the residue boundaries indicated. Parts that could be built reliably into the map are shown in a darker colour. **d**, Models of the four protein components forming the core-complex, EccB_5_, EccC_5_ EccD_5_ and EccE_5_. **e**, Representative region of density for each of the components visible in the local refined map for selected helices (threshold 0.22). For EccC_5_ region 26-61 density from the C6 map has been shown to highlight the connectivity between the TMH2 and the stalk domain.

**Extended Data Figure 5.**
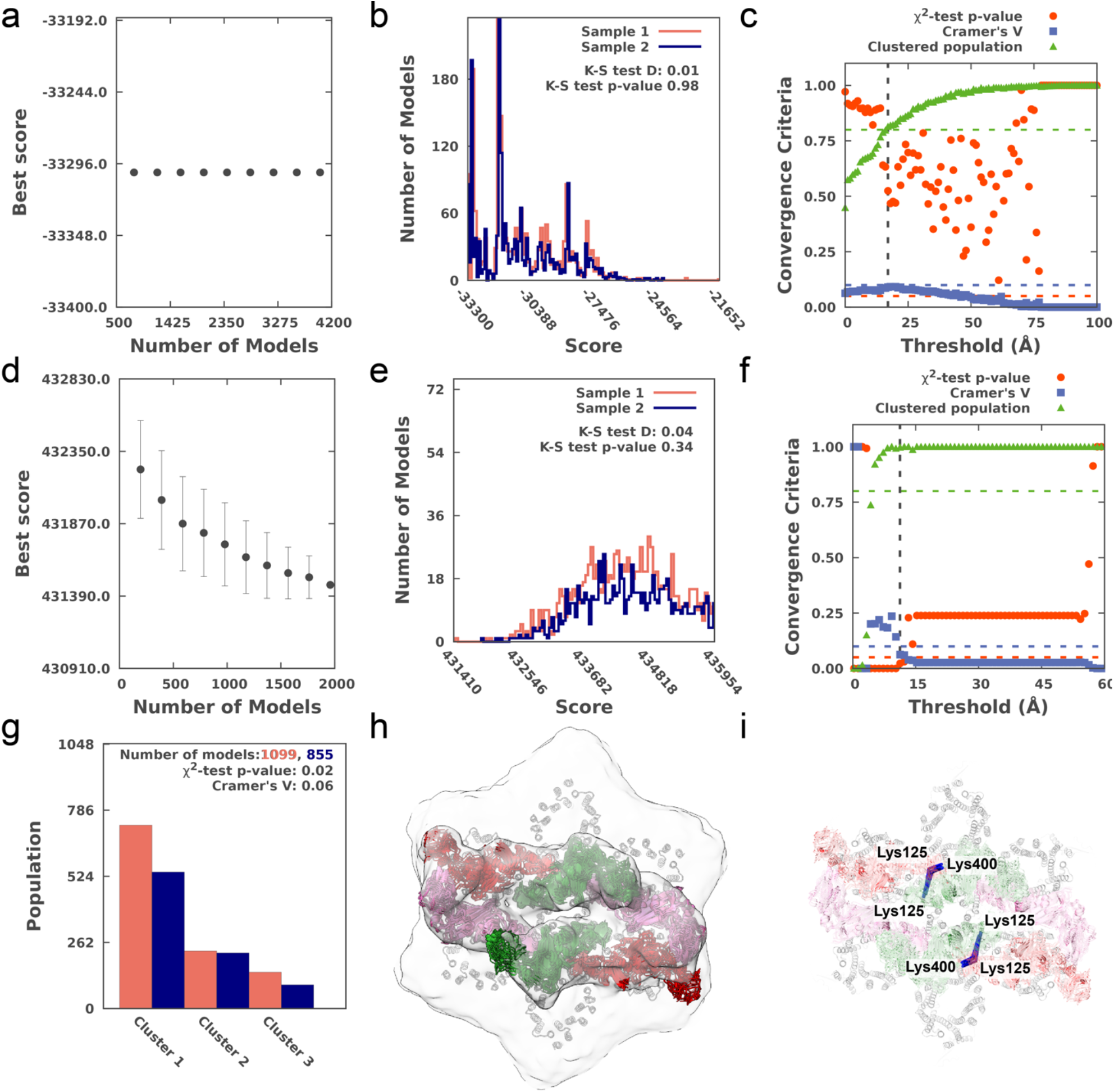
Assessment of the integrative model of the EccB periplasmic domain. **a**, Convergence of the first modeling stage of simultaneous fitting based on pre-calculated fits. The scores do not improve when more models are added indicating convergence. **b**, The score distributions of two independent samples of models are not statistically different also indicating convergence (p-value from the Kolmogorov-Smirnov two-sample test > 0.05, Kolmogorov-Smirnov two-sample test statistic, D < 0.3) **c**, Estimation of the sampling precision according to three criteria (y axis) of homogeneity of proportions in clustered models calculated for over increasing RMSD thresholds (x axis). The vertical dashed black line indicates the RMSD threshold at which the three conditions are satisfied (p-value > 0.05, Cramer’s V <0.10 (blue, horizontal dashed line) and the population of clustered structures >0.80 (green, horizontal dashed line), thus defining the sampling precision of 17 Å. **d, e, f**, Convergence of the refinement stage of the modeling tested as in a,b,c, defining the sampling precision after refinement of 11 Å. Note that the scores have a different scale as they are calculated differently in the refinement stage. **g**, Population of models from the two independent samples in the three clusters obtained by clustering at an RMSD threshold of 7 Å. **h**, Top ten scoring models of the biggest and the top cluster shown within the filtered EM map as used for modeling. The two-fold symmetrical pairs of EccB are shown in red, pink and green colours. These models are also the top scoring models out of all constructed models. Models from the two remaining clusters (not shown) are similar but with domain swaps of neighbouring EccB_5_s and thus having worse connectivity and total scores. The precision of the cluster 1 is 7 Å, which defines the overall precision of the EccB model. **i**, The two crosslinks within EccB_5_ (Lys125-Lys125 and Lys125-Lys400, blue bars) are satisfied by the models (distance < 30 Å).

**Extended Data Figure 6.**
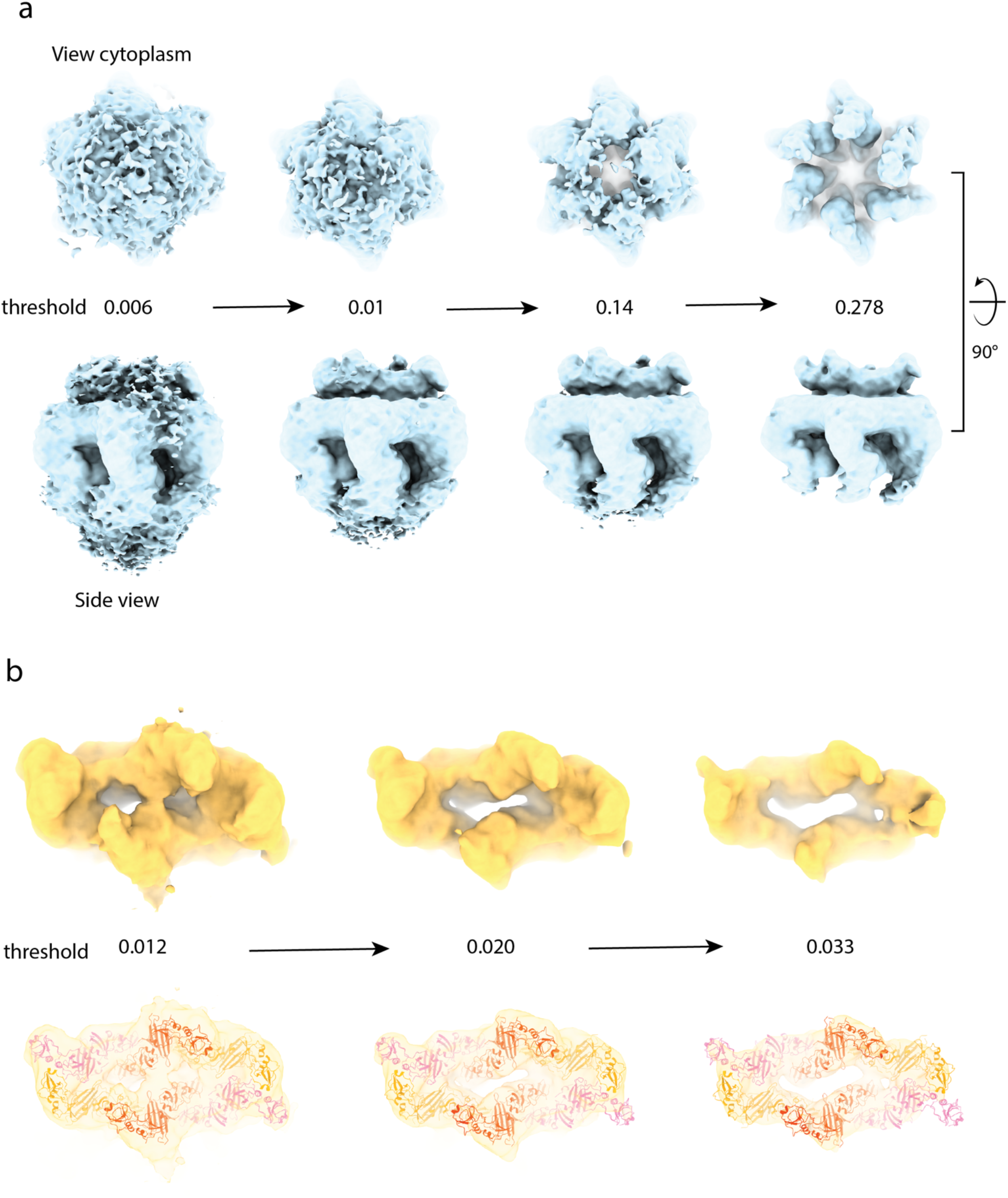
Low resolution features in the ESX-5 map. **a**, To visualise the density corresponding to the three EccC_5_ ATPase domains the C1 map has been displayed at different thresholds, with a view from the cytoplasmic side in the upper panel and from the side in the lower panel. At lower thresholds density corresponding to the cytoplasmic domains of EccC_5_ becomes visible. **b**, Visualising the EccB_5_ cleft, the density on the periplasmic side of the complex present in the C1 map has been shown at different thresholds. The cleft leads to the section pore in the membrane creating a channel. **c**, Highest scoring integrative model of periplasmic EccB_5_ fitted into the EM map, displayed at different thresholds.

**Extended Data Figure 7.**
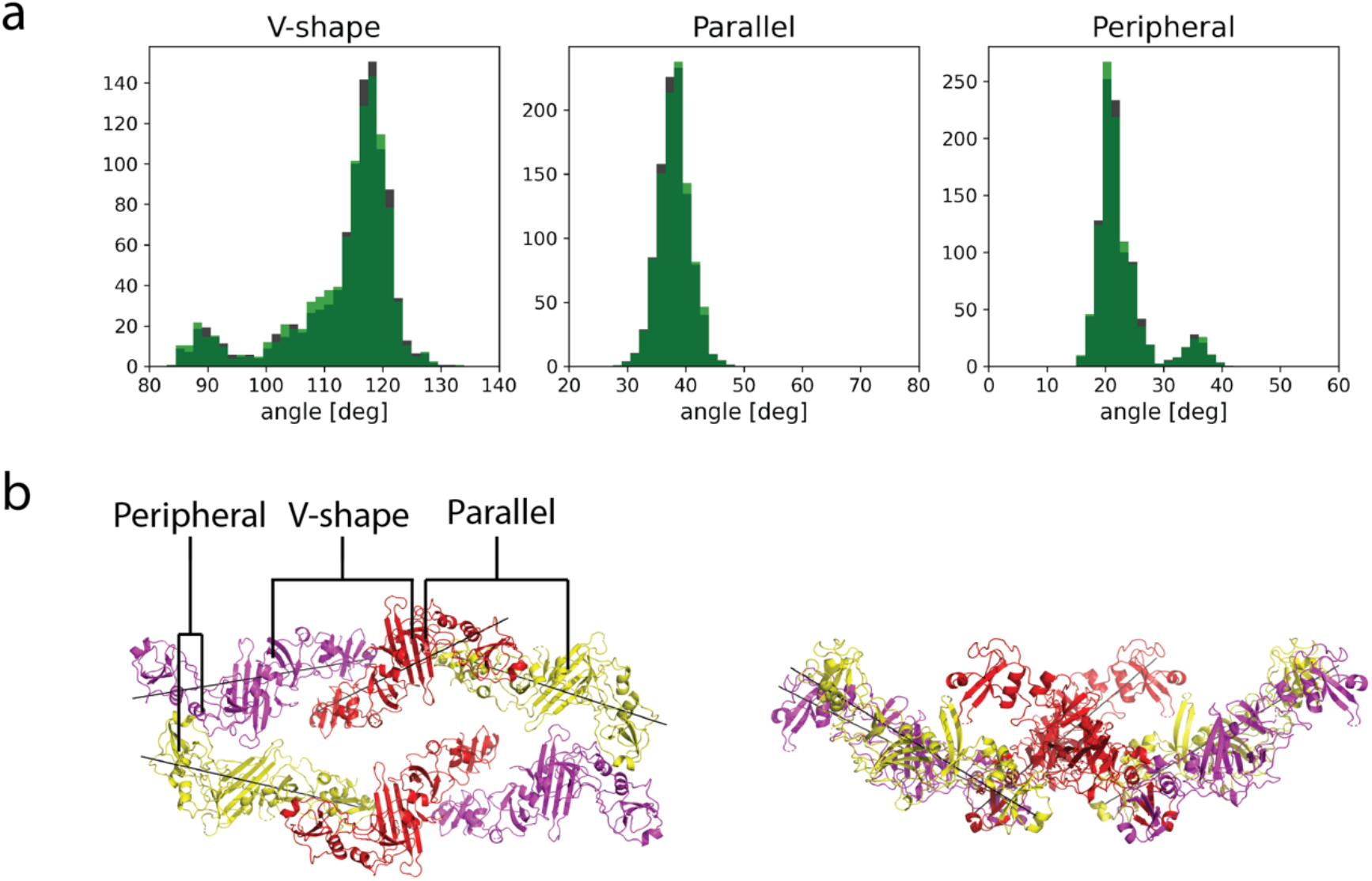
Analysis of angles between EccB dimers. **a**, Distribution of angles between three different EccB_5_ dimers in 1286 models obtained by integrative modelling. The distributions for symmetry related dimers are shown in green and grey. **b**, Angles between the dimers were measured using best-fit lines to the chain model coordinates determined using Principal Component Analysis. The lines for a representative model of the periplasmic EccB domains are shown in black.

**Extended Data Table 1.**
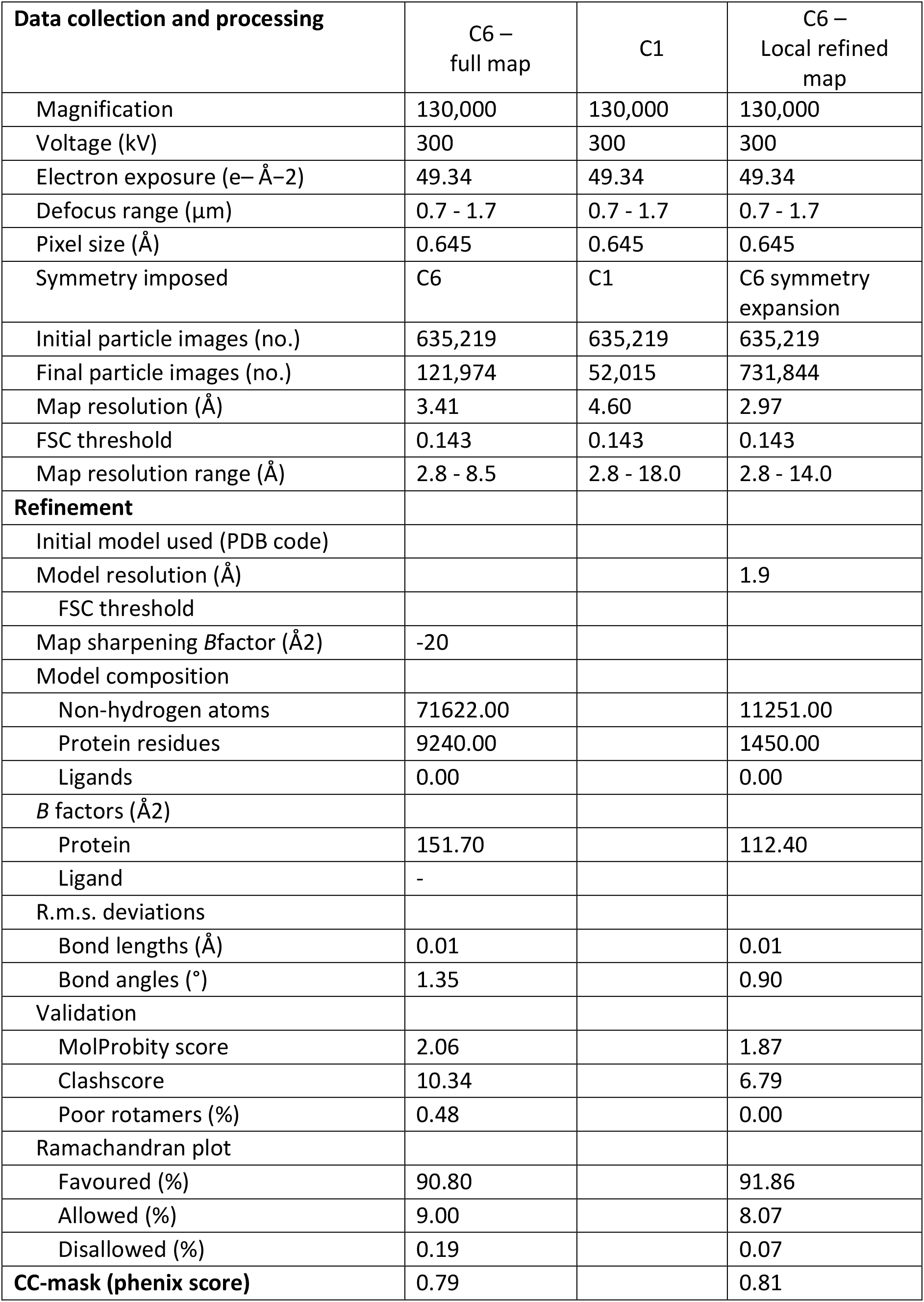
Cryo-EM and model building statistics.

| **Data collection and processing** | **C6 –**<br>**full map** | **C1** | **C6 –**<br>**Local refined**<br>**map** |
| --- | --- | --- | --- |
| Magnification | 130,000 | 130,000 | 130,000 |
| Voltage (kV) | 300 | 300 | 300 |
| Electron exposure (e<sup>-</sup> Å<sup>-2</sup>) | 49.34 | 49.34 | 49.34 |
| Defocus range (μm) | 0.7 - 1.7 | 0.7 - 1.7 | 0.7 - 1.7 |
| Pixel size (Å) | 0.645 | 0.645 | 0.645 |
| Symmetry imposed | C6 | C1 | C6 symmetry<br>expansion |
| Initial particle images (no.) | 635,219 | 635,219 | 635,219 |
| Final particle images (no.) | 121,974 | 52,015 | 731,844 |
| Map resolution (Å) | 3.41 | 4.60 | 2.97 |
| FSC threshold | 0.143 | 0.143 | 0.143 |
| Map resolution range (Å) | 2.8 - 8.5 | 2.8 - 18.0 | 2.8 - 14.0 |
| **Refinement** |  |  |  |
| Initial model used (PDB code) |  |  |  |
| Model resolution (Å) |  |  | 1.9 |
| FSC threshold |  |  |  |
| Map sharpening *B*factor (Å<sup>2</sup>) | -20 |  |  |
| Model composition |  |  |  |
| Non-hydrogen atoms | 71622.00 |  | 11251.00 |
| Protein residues | 9240.00 |  | 1450.00 |
| Ligands | 0.00 |  | 0.00 |
| *B* factors (Å<sup>2</sup>) |  |  |  |
| Protein | 151.70 |  | 112.40 |
| Ligand | - |  |  |
| R.m.s. deviations |  |  |  |
| Bond lengths (Å) | 0.01 |  | 0.01 |
| Bond angles (°) | 1.35 |  | 0.90 |
| Validation |  |  |  |
| MolProbity score | 2.06 |  | 1.87 |
| Clashscore | 10.34 |  | 6.79 |
| Poor rotamers (%) | 0.48 |  | 0.00 |
| Ramachandran plot |  |  |  |
| Favoured (%) | 90.80 |  | 91.86 |
| Allowed (%) | 9.00 |  | 8.07 |
| Disallowed (%) | 0.19 |  | 0.07 |
| **CC-mask (phenix score)** | 0.79 |  | 0.81 |

**Extended Data Table 2.** Data collection and refinement statistics.

|  | **EccD<sub>5</sub><sup>129</sup> (PDB ID : )** |
| --- | --- |
| **Wavelength** | 0.9762 |
| **Resolution range** | 30.59 - 1.66 (1.719 - 1.66) |
| **Space group** | P 43 21 2 |
| **Unit cell** | 43.979 43.979 85.146 90 90 90 |
| **Total reflections** | 62821 (6520) |
| **Unique reflections** | 10348 (1011) |
| **Multiplicity** | 6.1 (6.4) |
| **Completeness (%)** | 98.92 (99.41) |
| **Mean I/sigma(I)** | 14.12 (1.04) |
| **Wilson B-factor** | 31.22 |
| **R-merge** | 0.04768 (1.486) |
| **R-meas** | 0.05195 (1.61) |
| **R-pim** | 0.01979 (0.5997) |
| **CC1/2** | 0.999 (0.451) |
| **CC\*** | 1 (0.788) |
| **Reflections used in refinement** | 10346 (1011) |
| **Reflections used for R-free** | 521 (48) |
| **R-work** | 0.2280 (0.3374) |
| **R-free** | 0.18 (0.2272) |
| **CC(work)** | 0.956 (0.647) |
| **CC(free)** | 0.933 (0.510) |
| **Number of non-hydrogen atoms** | 740 |
| **macromolecules** | 701 |
| **solvent** | 39 |
| **Protein residues** | 91 |
| **RMS(bonds)** | 0.007 |
| **RMS(angles)** | 1.21 |
| **Ramachandran favoured (%)** | 100.00 |
| **Ramachandran allowed (%)** | 0.00 |
| **Ramachandran outliers (%)** | 0.00 |
| **Rotamer outliers (%)** | 0.00 |
| **Clashscore** | 0.70 |
| **Average B-factor** | 25.91 |
| **macromolecules** | 25.16 |
| **solvent** | 39.37 |
| **Number of TLS groups** | 1 |

Statistics for the highest-resolution shell are shown in parentheses.

